# Signaling and Mechanics influence the number and size of epithelial rosettes in the migrating zebrafish Posterior Lateral Line primordium

**DOI:** 10.1101/2025.05.17.650326

**Authors:** Abhishek Mukherjee, Michael Hilzendeger, Arin Rinvelt, Sana Fatma, Megan Schupp, Damian Dalle Nogare, Ajay B Chitnis

## Abstract

A prepattern of Fgf signaling triggers formation of epithelial rosettes as protoneuromasts form periodically in the migrating Posterior Lateral Line primordium. However, the number and size of epithelial rosettes is influenced by the balance of mechanical interactions that promote or oppose their formation. Selective slowing of leading cells in the primordium can result in the fusion of two rosettes to form one larger one, while slowing of trailing cells can result in splitting of a previously formed rosette to form two smaller ones. These observations can be accounted for by mechanics-based models, where local interactions associated with apical constriction and cell adhesion promote formation of rosettes, while tension along the length of the primordium, influenced by the relative efficacy of leading and trailing cell migration, opposes their formation. We describe computational models that illustrate how the relative speed of leading versus trailing cells, as well as changes in cell adhesion and mechanical coupling, can influence the pattern of protoneuromast formation and deposition by the migrating primordium. Our studies illustrate how signaling and mechanics together influence morphogenesis in the migrating primordium.

## Introduction

The development of an embryo represents a myriad of self-organizing processes resulting from interactions at multiple scales to determine robust emergence of fate, form and function in the growing animal. Conceptual frameworks developed to account for development often emphasize interactions operating at specific scales or via specific cellular, molecular, genetic and physical mechanisms. The challenge has been to develop an integrated view that recognizes their collective and interactive contribution. The relative simplicity and accessibility of the zebrafish posterior lateral line system for live imaging and for cellular, molecular and genetic manipulation has led to its emergence as an attractive system to understand development in a more integrative manner (Ghysen and Dambly-Chaudiere 2007).

The zebrafish posterior lateral line primordium is a group of about 140 cells (Nogare, Nikaido et al. 2017) that migrates under the skin, over the horizontal myoseptum, from near the ear to the tip of the tail, periodically forming and depositing sensory organs called neuromasts. These neuromasts spearhead formation of the Posterior Lateral Line, a sensory system that evolved to allow fish and amphibians detect the pattern of water follow over their body surface (Ghysen and Dambly-Chaudiere 2007). Each neuromast has sensory hair cells at its center, which are surrounded by support and mantle cells that cluster to form an epithelial rosette. Nascent neuromasts, or “protoneuromasts,” are periodically formed within the migrating primordium as interactions between primordium cells determine the specification of a sensory hair cell progenitor (SHCP) at the center of each protoneuromast. Each central sensory hair cell progenitor is surrounded by epithelialized cells that apically constrict and are recruited to form epithelial rosettes. Formation of protoneuromasts starts at the trailing end of the migrating primordium, and as they mature, new protoneuromasts are sequentially formed progressively closer to the leading end of the primordium. Eventually, trailing cells lose their capacity for migration and are deposited as neuromasts if they were successfully incorporated into protoneuromasts, or as inter-neuromast cells between the periodically deposited neuromasts, if not.

The sequential formation of protoneuromasts and collective migration of the primordium along the horizontal myoseptum is coordinated by polarized Wnt and FGF signaling in the primordium (Aman and Piotrowski 2008). Wnt activity dominates at the leading end, while it is relatively low at the trailing end. In the context of this polarized activity, Wnt signaling promotes expression of two FGF ligands, Fgf3 and Fgf10. However, Wnt active cells also express factors that inhibit FGF signaling and this prevents them from responding to the FGF ligands they produce (Aman and Piotrowski 2008, Matsuda, Nogare et al. 2013). Instead, these FGFs initiate FGF signaling at a distance, at the trailing end, where there is less inhibition of FGF signaling by Wnt activity. FGF signaling in this trailing domain determines expression of the diffusible Wnt inhibitor Dkk1b, where, together with other factors, it inhibits Wnt activity. By inhibiting FGF activity and by promoting expression of factors that promote Wnt activity, Wnt active cells, indirectly and directly promote Wnt activity. The *local activation* of Wnt signaling, coupled with its *long-range inhibition* by FGF signaling, constitutes a patterning system that has the potential to initiate FGF signaling in a center-biased pattern (Dalle Nogare and Chitnis 2020). This center-biased FGF signaling center initiates formation of a nascent protoneuromast by coordinating both the specification of a central *atoh1a*-expressing SHCP and reorganization of its surrounding cells to form an epithelial rosette. While Wnt activity inhibits FGF signaling and morphogenesis of protoneuromasts, FGF signaling promotes epithelialization, expression of factors like *shroom3* that promote apical constriction, and reorganization of cells to form epithelial rosettes (Ernst, Liu et al. 2012). Center-biased FGF signaling also determines specification of a central *atoh1a*-expressing SHCP, which subsequently contributes to maturation of the protoneuromast by stabilizing the reorganization of surrounding cells as epithelial rosettes (Lecaudey, Cakan-Akdogan et al. 2008, Matsuda and Chitnis 2010, Kozlovskaja-Gumbriene, Yi et al. 2017). Maturation of the trailing protoneuromast also contributes to inhibition of Wnt signaling in the trailing zone. This restricts Wnt activity to a smaller leading zone, establishing the conditions for the emergence of another center-biased FGF signaling domain in the wake of the shrinking Wnt system and the formation of a new protoneuromast. With each cycle, new protoneuromasts form progressively closer to the leading end.

FGFs secreted by leading Wnt active cells also serve as directional migratory cues for trailing cells (Dalle Nogare, Somers et al. 2014). As FGF-responsive trailing cells migrate toward the leading cells, the caudal migration of the primordium follows a path defined by the expression of the chemokine Cxcl12a (previously known as sdf1a) (Li, Shirabe et al. 2004, Sapede, Rossel et al. 2005, Haas and Gilmour 2006), guided by a self-generated local gradient of chemokine activity. Wnt activity in the primordium determines polarized expression of two chemokine receptors, chemokine (C-X-C motif) receptor 4b (Cxcr4b) and atypical chemokine receptor 3b (Ackr3b, previously Cxcr7b) (Haas and Gilmour 2006, Dambly-Chaudiere, Cubedo et al. 2007, Valentin, Haas et al. 2007). Wnt activity promotes *cxcr4b* expression in a leading zone, while it inhibits expression of *ackr3b,* resulting in the restriction of its expression to a trailing zone, where Wnt signaling is low (Aman and Piotrowski 2008). Ackr3b-expressing cells in the trailing zone internalize Cxcl12a and degrade it to determine formation of a local Cxcl12a gradient (Dona, Barry et al. 2013, Venkiteswaran, Lewellis et al. 2013, Bussmann and Raz 2015). Cxcr4b-expressing cells in the leading zone respond to this self-generated gradient to determine directed migration of the primordium. In this manner, chemokine and FGF signaling in leading and trailing cells, respectively, play complementary roles in coordinating the primordium’s directed migration.

One consequence of leading and trailing cells migrating in response to different mechanisms is that mechanical tension along the length of the primordium, associated with collective migration, is influenced by the relative efficiency of migration determined by these two signaling systems. Though a prepattern of FGF signaling seeds periodic formation of protoneuromasts, this study shows that morphogenesis of the epithelial rosettes is initially unstable, and that the number and size of the rosettes can be altered, at least transiently, by manipulations that potentially alter mechanical tension along the length of the primordium. Selective slowing of leading cells in the primordium can result in the fusion of two rosettes to form one larger one, while slowing of trailing cells can result in splitting of a previously formed rosette to form two smaller ones. We describe the development of two classes of computational models that illustrate how the local interactions associated with apical constriction and cell adhesion could promote clustering and formation of rosettes, while tension along the length of the primordium could serve as a long-range force that opposes such aggregation. In this manner, the models illustrate how a mechanical version of *local activation* and *long-range inhibition* could influence the pattern of cell clustering along the length of the primordium. We suggest that while FGF signaling creates a prepattern that coordinates the formation of protoneuromasts with a SHCP at the center of each epithelial rosette, tissue self-organization also allows for spontaneous periodic cell clustering because of mechanical interactions. Signaling and mechanics collectively determine the pattern of cell clustering in the migrating primordium.

## Results

### Manipulation of *cxcl12a* chemokine signaling slows leading cells in the PLLp

Interference with chemokine-dependent migration can be achieved by using heat shock to induce exaggerated and broad *cxcl12a* expression or by partially knocking down *cxcl12a* with morpholinos. To exaggerate *cxcl12a* expression, *Tg(cldnb:lyn-egfp); TgBAC(cxcr4b:h2a-mcherry); Tg(hsp:sdf1a)* triple transgenic embryos were heat-shocked between 27-28 hpf (hours post fertilization) at 37.5°C for 20 minutes. As shown previously (Dalle Nogare, Somers et al. 2014), broad induction of *cxcl12a* expression initially slows cells in the leading domain of the primordium before primordium migration eventually stalls relative to the wildtype control (Figure 1A, Movies S1 and S2), as induced protrusive activity in all directions interferes with directed migration of the primordium. Eventually, migration of the primordium stalls for approximately 7-8 hours before resuming in the caudal direction (Movie S2) (Lau, Feitzinger et al. 2020, Wong, Newton et al. 2020).

**Figure 1:**
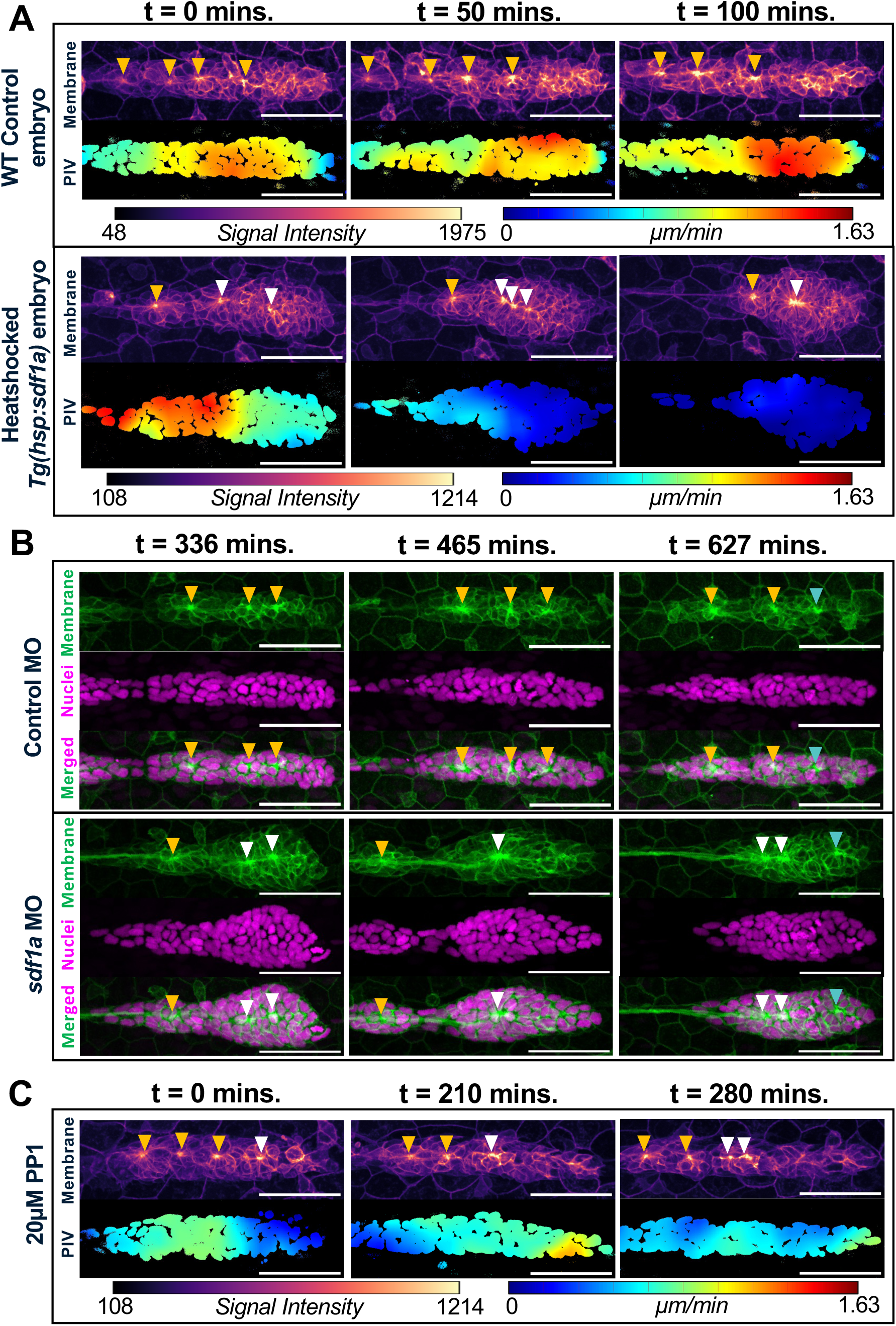
**(A)** Particle Image Velocimetry (PIV) maps, with warm hues indicating relatively high cell velocities (µm/min) in the horizontal direction, of primordia overlaid on cell nuclei shown over time, with a WT Control PLLp (top panel) compared to a heat shocked *hsp:sdf1a* PLLp (bottom panel). Respective membrane images are shown with an intensity-based hue (warm = high signal intensity) to clearly visualize apical constrictions for reference. Corresponding color bars are shown below the respective panels. **(B)** PLLps of Control morpholino injected (top panel) and *sdf1a* morpholino injected (bottom panel) embryos with membrane (green), nuclei (magenta), and merged channels shown over time to depict fusion of apical constrictions in the experimental set. **(C)** PIV maps of 20µm PP1 drug treated PLLp with their corresponding membrane images (warm hues indicating relatively high cell velocities (µm/min) in the horizontal direction or high signal intensity respectively) across time. **(A-C)** Yellow arrowheads indicate pre-existing constrictions that do not fuse over time. White arrowheads indicate constrictions that fuse or split over time. Cyan arrowheads denote new constrictions that appear in the leading domain of the PLLp. Corresponding color bars are shown below the respective panels. Note that deposited neuromasts are shifted out of the frame of view.

Time lapse imaging of the primordium coupled with analysis of cell movement with Particle Image Velocimetry (PIV) showed that, following heat shock-induced overexpression of *cxcl12a*, leading cells initially moved slower than trailing cells (Figure 1A; Movies S3, S4). The relative slowing of leading cells coupled with continued movement of trailing cells was accompanied by some apical constrictions moving closer to each other and eventually fusing to form a single larger rosette.

Interference with chemokine-dependent migration of cells in the leading zone of the primordium was also achieved with partial knock-down of *cxcl12a*. Injection of 2.0 ng of *cxcl12a* MO did not eliminate *cxcl12a*-dependent migration but rather, it appeared to create stretches with low enough *cxcl12a* to transiently slow or stall migration of primordium (Figure 1B; Supplementary Figure S1A-B; Movies S5, S6). Though this manipulation had an opposite effect on *cxcl12a* levels, effects on epithelial rosette formation were like those observed with over-expression. That is, the leading cells stalled or slowed down briefly leading to fusion of some apical constrictions. Similar effects suggested that reorganization of apical constrictions did not correlate with changes in the level of chemokine signaling, but rather, to their common effect of selectively slowing migration of leading cells in the primordium, while trailing cells continue to move, resulting in concertina-like compression of the migrating primordium.

#### Inconsistent coupling of *atoh1a* expression and apical constrictions in nascent protoneuromasts

In each developing protoneuromast, a central *atoh1a*-expressing SHCP is expected to be associated with surrounding cells that reorganize to form an epithelial rosette, which can be identified by its apical constriction (Figure 2A). Examination of *atoh1a* expression before heat shock and 4 hours after heat shock-induced *cxcl12a* expression, showed that the number of *atoh1a*-expressing cells occasionally exceeded the number of detected apical constrictions (Figure 2A,B). As a single *atoh1a*-expressing cell is expected to be associated with each protoneuromast and its corresponding epithelial rosette, observing multiple *atoh1a-*expressing cells associated with a single apical constriction is consistent with fusion of initially adjacent protoneuromasts, resulting in the formation of a single larger epithelial rosette, as observed in timelapse images.

**Figure 2:**
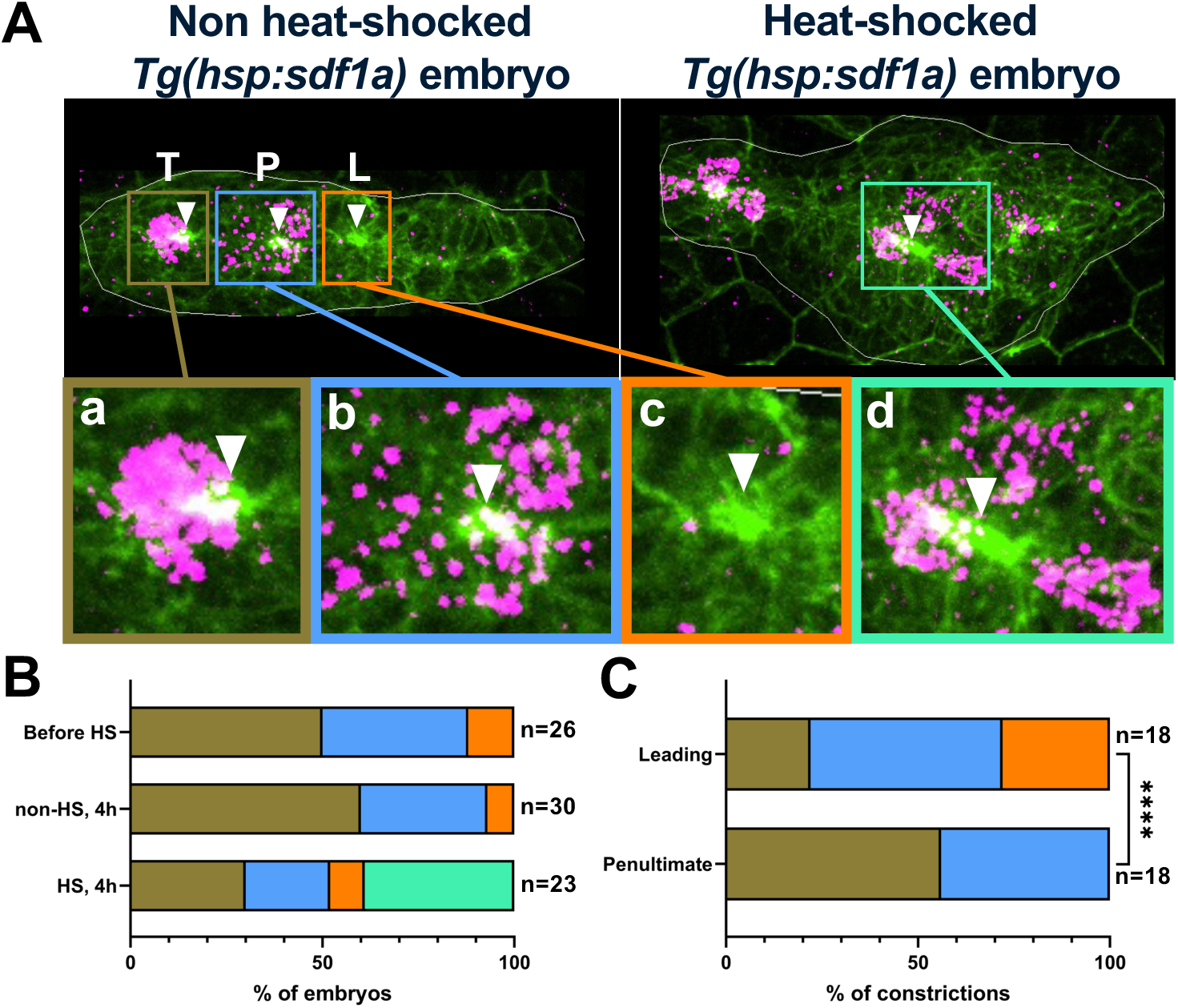
**(A)** Representative (top) images demonstrating the various types of observed patterns associating apical constrictions with *atoh1a* expression. T, P, and L denote trailing, penultimate, and leading protoneuromasts respectively. Membrane is shown in green, and RNA *in situ* hybridization for *atoh1a* is shown in magenta. White boundaries mark the outlines of the PLLps. **(a-d)** Magnified images of the regions of interest within colored rectangles in **(A)** showing 4 distinct phenotypes observed experimentally**. (a)** (brown) = constriction associated with 1 *atoh1a* cell; **(b)** (blue) = constriction associated with diffused *atoh1a* expression; **(c)** (orange) = constriction having no associated *atoh1a* expression; **(d)** (green) = constriction associated with more than 1 *atoh1a* cells). **(B)** Contingency plot depicting the 4 observed phenotypes in **(A)**, scored for three experimental conditions, and presented as percentages of the total number of embryos. **(C)** Contingency plot scoring the number of leading (L) and penultimate (P) constrictions showing the 4 observed phenotypes in **(A)**, within all non-heat-shocked *Tg(hsp:sdf1a)* embryos. Asterisks indicate significance by Fisher’s exact test (**** p < 0.0001). Total number of embryos **(B)** and total number of constrictions **(C)** are noted on the right of the respective bars.

However, examination of the relationship between the position of apical constrictions and the underlying *atoh1a* expression pattern revealed that apical constrictions were often not associated with a well-defined single *atoh1a*-expressing cell. Instead, they were associated with a diffuse pattern of *atoh1a*, especially in nascent protoneuromasts in the leading zone (Figures 2). In some cases, the apical constriction preceded the appearance of *atoh1a* expression in the nascent protoneuromast (Figure 2A-c). This is consistent with mechanisms that determine specification of a central *atoh1a* expressing cell and reorganization of cells to form epithelial rosettes operating independently and in parallel when protoneuromast formation is initiated. As protoneuromasts mature, the relationship between the central *atoh1a*-expressing cells and the surrounding cells forming the epithelial rosette becomes increasingly well-defined (Figure 2C). In some cases, diffuse or poor *atoh1a* expression was also seen in the most trailing protoneuromast prior to its deposition. However, this is likely to be associated with the *atoh1a*-expressing SHCP in the most mature trailing protoneuromast dividing to form two sensory hair cells prior to deposition, during which time *atoh1a* expression is lost. As a result, trailing protoneuromasts were not included in our analysis.

#### Inhibition of Src kinase signaling selectively slows trailing cells

PP1, a Src kinase inhibitor, had previously been used to inhibit collective migration of the primordium (Neelathi, Dalle Nogare et al. 2018). Time lapse imaging data coupled with PIV analysis showed that as primordium migration slows in response to 20µM PP1, trailing cells begin to slow earlier than leading cells (Figure 1C, t = 210 mins.), before the entire primordium eventually slows down (Figure 1C, t = 280 mins., Supp. Movie S7). In this context, selective slowing of trailing cells versus the leading cells causes the primordium to stretch and this tension is accompanied by splitting of some preexisting apical constrictions to form two (Figure 1C, t = 280 mins.).

#### Models to account for dynamic reorganization of apical constrictions

The fusion or splitting of preexisting apical constrictions, when leading or trailing cells preferentially slow down, suggested that formation of epithelial rosettes is influenced by the tension along the length of the primordium, which is influenced by the relative efficacy with which leading and trailing cells migrate. When leading cells are slower than trailing cells, the primordium length decreases, and the tension along the length of the primordium, which opposes cell clustering, reduces. This allows preexisting apical constrictions to come together to form fewer larger rosettes. Conversely, when trailing cells selectively slow down and leading cells keep moving, the primordium lengthens, tension opposing cell clustering increases, and preexisting rosettes are pulled apart forming more and smaller rosettes. Two sets of models were developed to test this hypothesis, one an Agent Based Model developed with NetLogo and the other a Cellular Potts model developed with CompuCell3D.

#### An Agent-Based model of dynamic cell clustering in the primordium

The primordium in the Netlogo ABM was represented as a column of 150 *turtles*, 5 wide and 30 long, approximating the number of cells in the actual primordium at the start of its migration (Figures 3,4). Consistent with the Wnt active domain occupying the leading 60% of the primordium, **WNTers**, representing cells in the Wnt active domain were in the leading 60% of the model primordium, while the trailing 40% was occupied by **FGFers** (Figure 3A).

**Figure 3:**
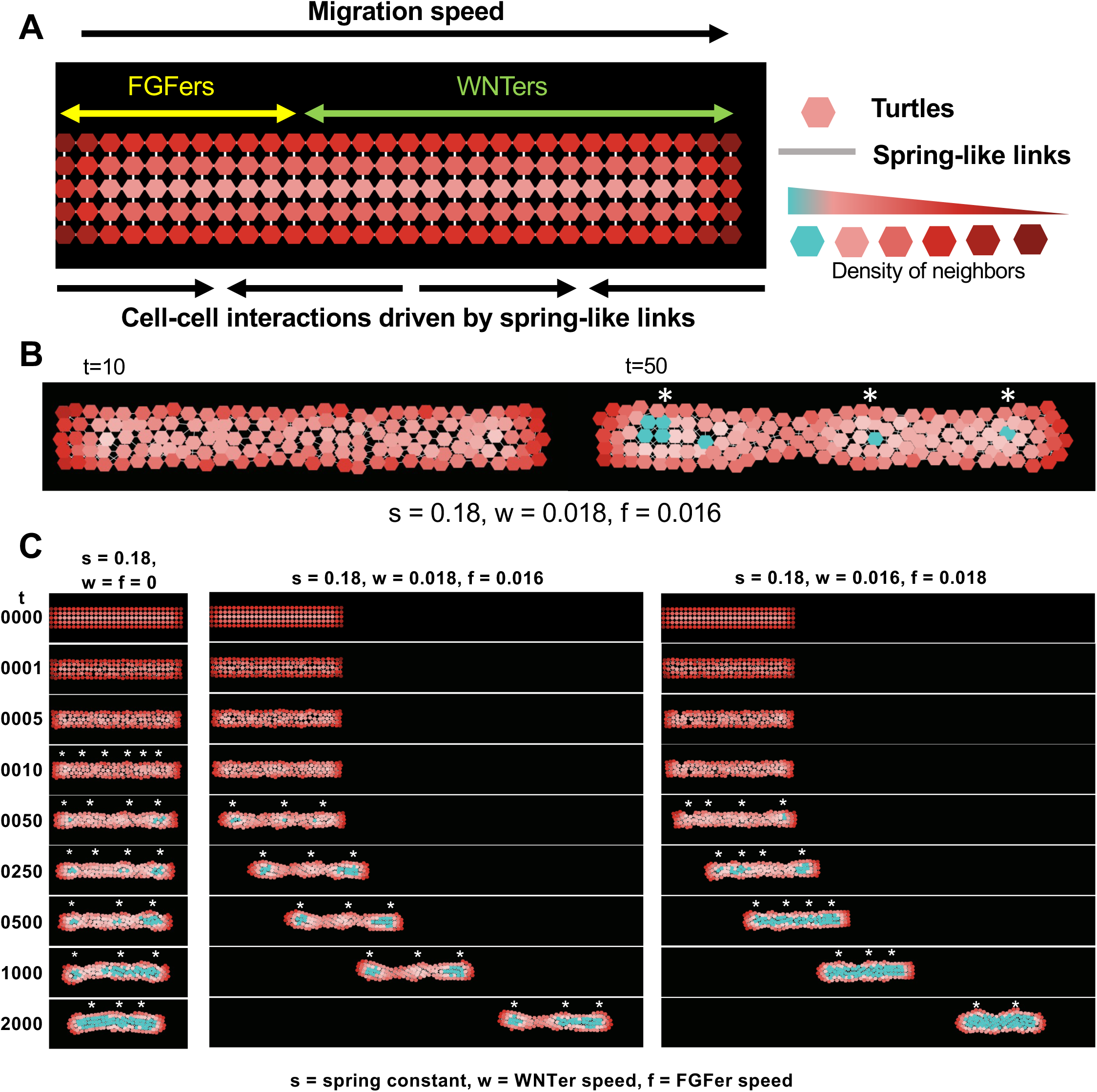
**(A)** Agent-based model setup with direction of migration is shown. Wnt domain extends up to 60% of the model PLLp length from the leading edge, and FGF domain occupies the rest of the 40% of the model PLLp length. Hexagon-shaped turtles (WNTers and FGFers) are connected to their neighbors with spring-like links and their hue changes with the number of neighbors, as shown in the legend. **(B)** An example simulation is shown with spring constant, s = 0.18, WNTer speed, w = 0.018, FGFer speed, f = 0.016. Due to cell-cell adhesive and repulsive interactions, cells reorganize to form local clusters (3 clusters shown here with cyan depicting high neighbor density). **(C)** Representative simulations without migration (left column), with WNTer speed > FGFer speed (middle column), and with WNTer speed < FGFer speed (right column).

**Figure 4:**
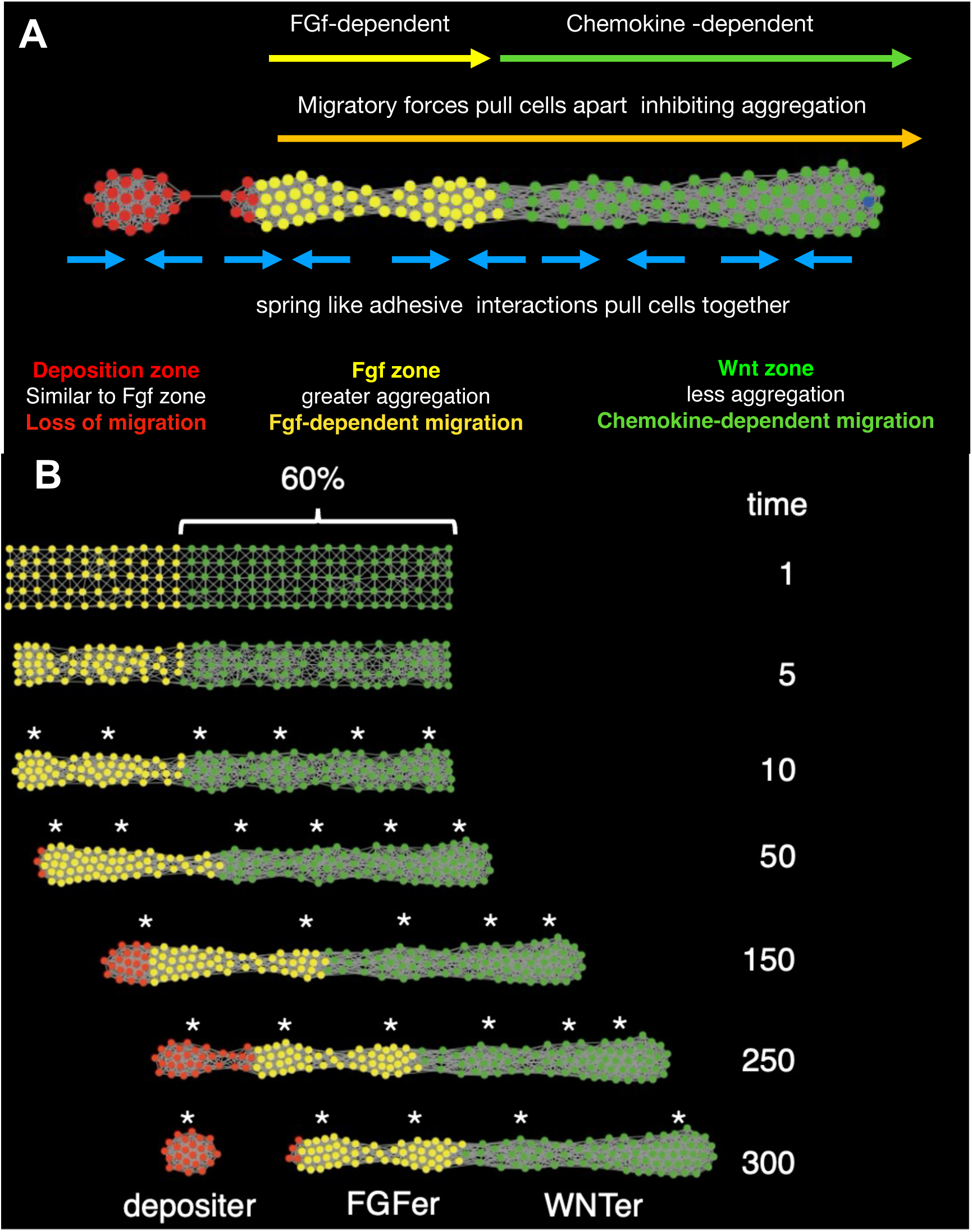
**(A)** An agent-based model simulating the reduction of Wnt domain over time and the consequent neuromast deposition. All agents are represented as circles. WNTers (green) in the leading zone extend up to 60% of the PLLp length, followed by FGFers (yellow) making up the remaining 40%. Depositers (red) are the agents that are found beyond the PLLp length. The most critical behaviors that set these three types of agents apart from each other are highlighted. **(B)** Timed progression of the model PLLp shows that Depositers get deposited as the Wnt domain shrinks. Self-organized clusters are denoted with asterisks.

The model was used to visualize how spring-like interactions between cells can determine self-organized clustering of cells and how varying the “spring-constant” of *links* or the relative “speed” of **WNTers** and **FGFers** influences the pattern of cell aggregation. Aggregation of cells was visualized in two ways; first, the shade of red used for each turtle reflected how many neighbors it had within a defined radius (a brighter shade for more, dark shades for less). Second, when the number of turtles within a defined radius of a turtle exceeded a threshold, its color was set to cyan (Figure 3B). When **WNTers** and **FGFers** were assigned the same spring-constant (s) of 0.18 but no speed (w = f = 0) (Figure 3C (left panel) and Movie S8, top panel), turtles along the entire primordium began to form local clusters (white asterisks). Initially a larger number of smaller clusters formed, which, when allowed sufficient time, coalesced together to form fewer but larger aggregates. The aggregation of the turtles increased with increasing spring constant (Supp. Figure S2). When the speed of the leading **WNTers** was higher than the trailing **FGFers** (Figure 3C (middle panel) and Movie S8, middle panel), the resulting tension along the length of the primordium stretched the primordium, creating three relatively stable clusters. However, as the **WNTers** in the leading 60% of the primordium move at the same speed, eventually the leading two clusters began to coalesce to form a larger dense cluster (compare t = 1000 and t = 2000). When **WNTers** had a slower speed than **FGFers** (w = 0.016 and f = 0.018), coalescing of cells to form aggregates was accelerated and two dense clusters were formed by t = 2000 (Figure 3C (right panel) and Movie S8, bottom panel). These results, taken together, suggest that while contractile spring-like forces can promote aggregation of cells, tension caused by differential migration speeds can pull them apart and inhibit aggregation. Our results illustrate that even with a simple model, cell clustering and morphogenesis can be modulated by the interplay of contractile and tensile forces.

During the migration of the primordium in the embryo, the Wnt system progressively shrinks, as new protoneuromasts form in its wake. However, as the Wnt system shrinks, the primordium shrinks as well, and its length scales with the length of the Wnt system (Dalle Nogare and Chitnis 2017). In the model, the Wnt domain could be set to shrink at a defined rate and the length of the model primordium was adjusted so that it was consistently 1.6 times the current length of Wnt domain (keeping the size of the Wnt domain consistently at 60% the length of the primordium) (Figure 4A-B). **FGFers** that extended in the trailing direction beyond the permitted length of the primordium at any time were considered to have lost their capacity to migrate and were re-specified as a third breed of *turtles* called **Depositers**. **Depositer** parameters matched those of **FGFers**, except they had zero migration speed, did not proliferate, and had low *link* break-thresholds, so that as the rest of the *turtles* in the model primordium kept moving, **Depositers** broke their *links* with the **FGFers** and were “deposited”. In this manner the model was used to visualize how cell clusters form and deposit in the wake of a progressively shrinking Wnt system (Figure 4B, Movie S9).

While the Netlogo ABM illustrates how spring-like cell connections and migration-associated tension can influence self-organization of clusters within the migrating primordium, it has several limitations. One important limitation is that each cell is represented as a point on a lattice; it has no real spatial dimension, so spring-like connections between turtles in our model are used to simulate interactions that are likely to arise from both cell-cell adhesion and apical constriction. To more explicitly model these distinct interactions, Cellular Potts Models (CPMs) (Glazier-Graner-Hogeweg models) using the CompuCell3D modelling environment (Swat et al (2012)) were developed. In a CPM, each cell is represented as a set of pixels that define the position, size/shape, and degree of contact with other cells in the model. The first step in the development of the model is to define the different types of cells that will be interacting, their initial configuration, and features of the environment in which the cells will operate. Each different cell type has unique target parameters, including size (perimeter, surface area), degree of adhesivity to other cells of the same or different type, and movement in response to potential chemotactic influences on the cell. At each iteration of the program, pixels from the set representing each cell are randomly added, subtracted, or relocated within or adjacent to the cell’s current pixel set. This results in growth, shrinkage, shape alteration, net movement of each cell, or variation in contact with adjacent cells. Whether this change in the cell pixel position is accepted or rejected at each iteration depends on an energy function that computes whether the resulting change moves the cell closer or further from predefined target parameters (Eq. (2), Methods). Deviation of each cell’s new parameters from its pre-defined target parameters is associated with an energy cost, and the sum of these energies is used to determine if moving the pixel is associated with a net reduction in the energy state of the cell; if the net energy is reduced, the change is accepted, if not it is either rejected, or accepted at a low probability (Eq. (3), Methods). Iterations in this process of randomly moving pixels associated with each cell, and accepting or rejecting the change based on whether it lowers net energy of the system, allows it to evolve over time. In this way, the model simulates the potential trajectory of the system as cells dynamically change size, shape, position, and association with neighboring cells.

#### A Cellular Potts Model (CPM) recapitulates rosette formation and emergent behavior of the primordium

Although the top-view ABM provided us with valuable insights into the dynamics of self-organization of tissues, the interplay of apical contractility in apicobasally polarized cells and their migration is still unclear. Therefore, we created a side-view Cellular Potts model. The lateral line primordium migrates under the skin, over the horizontal myoseptum, and is made of cells with distinct shapes and behaviors. Its migration and distribution of cells with distinct morphology is best seen when viewing the primordium from its side (Figure 5A)

**Figure 5:**
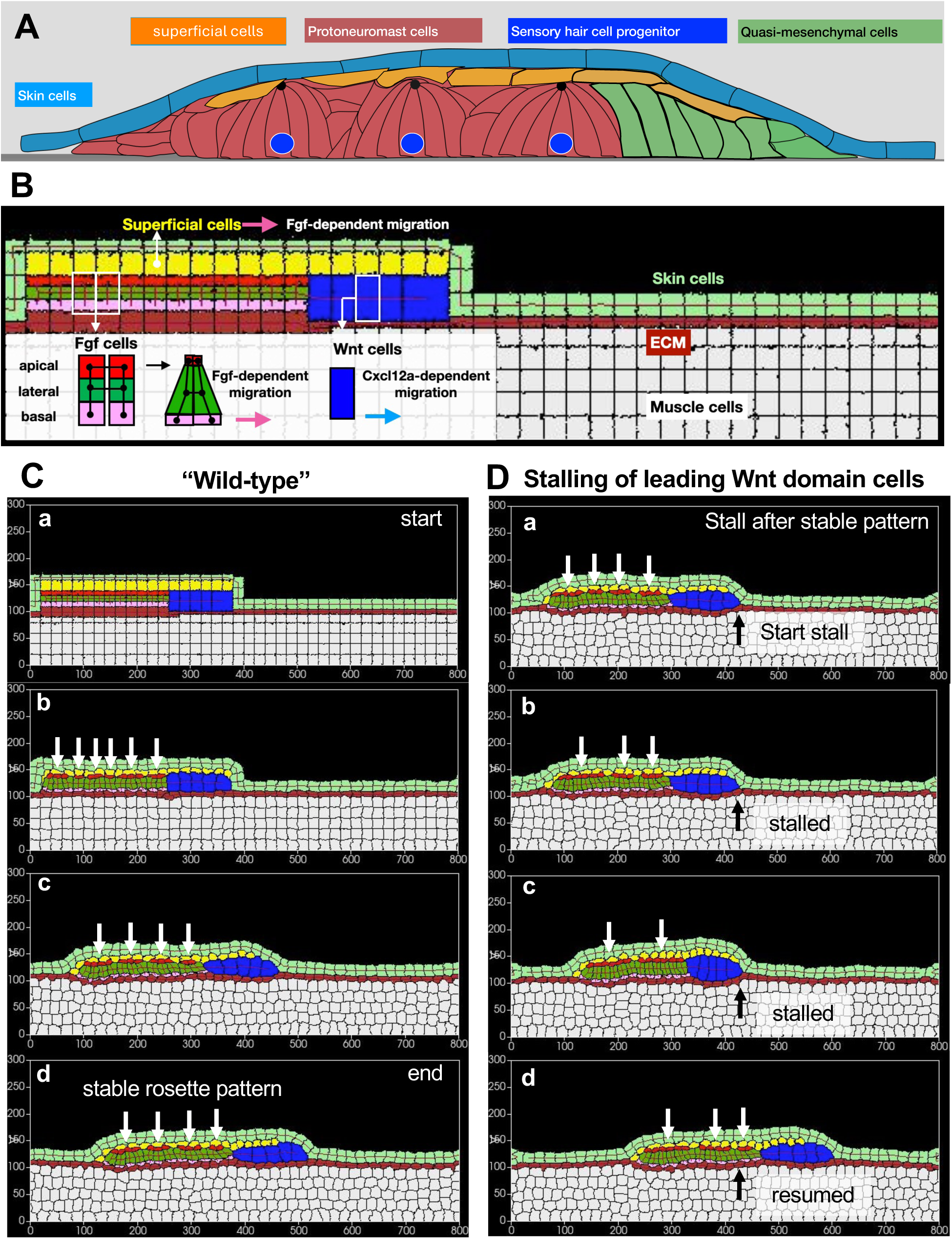
**(A)** A schematic showing the broad categories of cells that compose a PLLp and the environment it migrates through. **(B)** A schematic Cellular Potts model setup, incorporating different types of cells that make up the PLLp as well as the main components of their extracellular environment. Apicobasally polarized cells in the trailing Fgf domain are represented as composites of three compartments, apical (pink), lateral (green) and basal (pink) whose target parameters are specified individually. Spring-like *FocalPointPlasticity* (FPP) links shown in black. Increasing the spring-constant of the FPP link between adjacent apical compartments simulates apical constriction (black Arrow). Pink arrow represents a force that determines Fgf-dependent migration of the basal compartment of Fgf cells and superficial cells, blue arrow represents force associated with Cxcl12a-dependent migration of Wnt cells. **(C)** Progression of migration and rosette formation of a model wild-type PLLp through different stages, showing the start of a simulation **(a)**, initial re-organization of rosette-like structures as indicated by clustering of red apical domains **(b)**, and eventually locking into a stable rosette configuration**(c-d)**. **(D)** Progression of a model PLLp simulating stalling of the leading unpolarized blue cells, depicting start of the stalling event **(a)**, aggregation and reorganization of cells to form larger rosette-like structures in the stalled model **(b and c)**, and eventual splitting up of one of the rosettes to form smaller ones in **(d)**.

We modeled the cells in the leading domain (Wnt domain, blue) as not apicobasally polarized. The trailing domain is comprised of apicobasally-polarized FGF cells. Epithelialized FGF cells have apical, lateral, and basal compartments, each with unique behaviors and characteristics. While cells in this domain can form apical constrictions, they are adherent to each other along their entire length via membrane proteins and migrate with their basal lamellipodia. To simulate these distinctive regional properties in the CPM, polarized epithelial **FGF** cells were represented as a composite of three cell-types (**fgfapi** (red), **fgflat** (green), and **fgfbas** (pink)), each of which were assigned target parameters representing the unique features of the apical, lateral, and basal compartments, respectively. A relatively small target size for the **fgfapi** cell-type, coupled with short and strong spring-like *FocalPointPlasticity* links between apical compartments, helped simulate strong apical constriction in **FGF** cells (Appendix, Supplementary Tables 2-5).

Though direct information about the adhesive nature of each cell was not available, the following assumptions were made in the development of the CPM model. As there is no mixing of cells of one type with cells of another type in the primordium during migration, it was assumed that each distinct cell type was most adherent to cells of the same type. Second, as primordium cells move as a cohesive column, it was assumed that they adhere more to each other than to the overlying **skin** cells or the underlying **ECM** through which they migrate. Furthermore, the adhesion between **superficial** cells and the overlying **skin** cells, or that between the **fgfbas** compartment of **FGF** cells and underlying **ECM** cells, was set low enough to allow these primordium cells to move without getting persistently attached to these substrates.

Simulations using the CPM model, based on the assumptions described above, recapitulated many aspects of primordium behavior. First, the primordium cells migrated as a cohesive entity between the skin cells above and the ECM below. Second, the adhesivity of superficial cells to the overlying **skin,** and of the basal compartments of FGF cells to the underlying **ECM,** was high enough to keep them attached to these substrates, but low enough to allow detachment as the primordium moved forward.

Third, as the primordium cells began to migrate, adhesive and spring-like interactions between **FGF** cells determined the spontaneous clustering of cells to form “rosettes”. While initially some of the rosettes were unstable (Figure 5C(a-b)), the system settled into a stable configuration, shown in Figure 5C(c-d) and Movie S10, with four rosettes. Finally, these models enabled us to simulate experiments in which interference with Cxcl12a-dependent migration determined the slowing of leading cells and resulted in transient fusion of pre-existing, leading Wnt cells. This resulted in progressive fusion of preexisting four rosettes to first form three and then two larger rosettes (Figure 5D(a-c), Movie S11). Then, as migration of leading Wnt cells was restored, one of the fused rosettes split again to form two (Figure 5D(d)), recapitulating the phenomenon that had been seen in vivo. We further explored the influence of cell contractility and cell-cell adhesion in defining rosettes in the migrating system by increasing the contractility and adhesion strength of **fgflat** compartments (Appendix, Table 5). We observed that such a change led to the FGF cells to gradually aggregate such that initially formed smaller rosettes fuse to form a large rosette even when migration itself is not compromised (Figure S3, Movie S12), thereby overriding the tension in the column of cells normally caused by migration. This simulation reproduced a phenomenological observation from (Kozlovskaja-Gumbriene, Yi et al. 2017), where activating Notch across the entire primordium leads to the formation of a large rosette in the trailing domain.

## Discussion

The posterior Lateral Line primordium migrates under the skin, from near the ear to the tip of the tail, periodically forming and depositing neuromasts. The periodic and sequential formation of protoneuromasts within the migrating primordium requires the coordination of two patterning events; one involving cell fate specification, where an *atoh1a*-expressing cell becomes specified as a sensory hair cell progenitor at the center of each nascent protoneuromast, and the other involving morphogenesis, where surrounding cells reorganize to form an epithelial rosette. Previous studies have shown that FGF signaling initiates both these events; first, it promotes *atoh1a* expression, which gives cells the potential to become sensory hair cell progenitors, and second, FGF signaling promotes epithelialization of cells and the expression of *shroom3*, which promotes apical constriction and reorganization of cells to form epithelial rosettes.

The sequential periodic formation of protoneuromasts, starting from the trailing end of the primordium, is coordinated by polarized Wnt signaling. It dominates in the leading zone, where it inhibits FGF signaling and protoneuromast formation. At the same time Wnt signaling in leading cells determines expression of the FGFs that are delivered to trailing cells, where the activation of FGF signaling coordinates formation of nascent protoneuromasts. It was shown that previously described interactions between Wnt and FGF signaling define a reaction diffusion system, where local self-activation of Wnt signaling, coupled with long range inhibition of Wnt activity by FGF dependent Dkk1b expression, has the potential to determine self-organization of center-biased FGF signaling centers in the wake of a shrinking Wnt activity domain (Dalle Nogare and Chitnis 2020). These signaling centers create a prepattern of FGF signaling that initiates formation of protoneuromasts with their central *atoh1a*-expressing cell and surrounding cells organized to form epithelial rosettes. Once *atoh1a* expression becomes restricted to the central cell this cell becomes the source of signals that activate Fgf and Notch signaling in its neighbors, consolidating their organization as epithelial rosettes.

While the Fgf pre-pattern helps understand how specification of a central atoh1a-expressing cell becomes coupled to surrounding cells that form epithelial rosettes, our observations show that these events can become uncoupled. Manipulations that selectively slowed the leading cells allowed the column of primordium cells to compress along its length and become more rounded; this was accompanied by preexisting apical constrictions coming together to form fewer larger rosettes. As leading cells resumed migration, single constrictions sometimes split to form two once again, reflecting the relative plasticity of cell clustering. Similarly, when trailing cells were selectively slowed, the column of primordium cells stretched, and preexisting constrictions split to form two constrictions.

The uncoupling of mechanisms that determine specification of a central *atoh1a*-expressing cell from those that determine periodic formation of epithelial rosettes was illustrated by comparing *atoh1a* expression and position of apical constrictions. Though the uncoupling of *atoh1a*-expressing cells and apical constrictions was revealed in embryos with slowed leading cells, the relative independence of *atoh1a*-dependent sensory hair cell specification and epithelial rosette morphogenesis was suggested earlier by *atoh1a* knockdown experiments which showed that this does not prevent periodic epithelial rosette formation; the primordium continues to migrate, forming and depositing “neuromasts” that form relatively unstable epithelial rosettes (Lecaudey, Cakan-Akdogan et al. 2008). While Fgf normally initiates formation of epithelial rosettes, it has been shown that a prepattern of FGF signaling activity is not required for self-organization of multiple epithelial rosettes in the migrating primordium. In primordia in which there was broad induced expression of an activated form of Notch, without any pre-pattern, a few relatively large rosettes formed (Kozlovskaja-Gumbriene, Yi et al. 2017). Similarly, when FGF signaling was inhibited in control primordia, epithelial rosettes disassembled, and their formation was inhibited. However, when FGF signaling was inhibited in primordia with broad ectopic Notch activation, cells self-organized to form about three rosettes in the absence of any pre-pattern of Notch or FGF signaling (Kozlovskaja-Gumbriene, Yi et al. 2017). Together, these observations suggest that while FGF signaling may help initiate epithelial rosette formation, Notch activation is adequate to determine expression of factors that promote self-organization of primordium cells to form rosettes. This includes promotion of adhesion factors like Cdh-1 (E-cadherin) and EpCAM, along with tight junction components like CldnB and CldnE, which are expected to promote adhesion and apical constriction. These observations show that while a pre-pattern of individual FGF signaling centers may more reliably initiate periodic formation of protoneuromasts with their central *atoh1a*-expressing cells and associated epithelial rosettes, broad un-patterned expression of factors that promote adhesion and apical constriction is adequate to determine self-organization of cell clustering within the migrating primordium.

Agent based Models (ABMs) and Cellular Potts Models (CPMs) were used to illustrate how mechanical interactions mediated by cell adhesion and apical constriction could determine self-organization of cell clustering along the length of the primordium. The underlying principle in both types of models is that adhesive interactions coupled with apical constrictions pull cells together to promote cell clustering. Unimpeded, this process is self-amplifying, and the clusters can grow progressively larger. However, cell attachment to the substrate or migratory behavior that stretches the primordium generates tension along its length, opposing cell aggregation. Together, the local interactions that pull cells together to promote clustering, which are opposed via long range inhibition by tension, determine self-organization of cell clusters. Conceptually, this self-organizing process can be thought of as a mechanical version of the same principles of local activation and long-range inhibition associated with Wnt and FGF signaling described earlier that have the potential to determine self-organization of FGF signaling centers. The mechanical interactions that determine the spontaneous formation of cell clusters and their similarity to reaction-diffusion mechanisms described independently by Turing (Turing 1990) and Meinhardt (Gierer and Meinhardt 1972, Meinhardt 2012) had been described by Oster et al. (Oster, Murray et al. 1983) and by Harris et al. (Harris, Stopak et al. 1984) more than 50 years ago.

Previous analysis of signaling interactions mediated by the Wnt, FGF, and Notch signaling pathways has shown how they can determine self-organization of periodic FGF signaling centers with central *atoh1a*-expressing cells (Dalle Nogare and Chitnis 2020). Here, we have described how mechanical interactions between cells can also determine periodic cell clustering within the primordium, with both signaling and mechanical interactions capable of spontaneously generating these patterns without a pre-pattern. Nevertheless, in the migrating primordium, as the Wnt system shrinks, signaling and mechanical interactions that determine formation of center-biased FGF signaling simultaneously initiate *atoh1a* expression and clustering of cells, coupling the interactions required for the formation of a new protoneuromast. Although these patterning mechanisms can operate independently, they mutually amplify each other and eventually become synchronized. Nevertheless, morphogenesis of an epithelial rosette is not reliably associated with a central *atoh1a*-expressing cell in nascent protoneuromasts, and the relationship can be more easily uncoupled by manipulations that result in stretching or compression of the migrating primordium. As *atoh1a*-expression becomes self-sustaining in maturing protoneuromasts, the central sensory hair cell progenitor stabilizes as a signaling source, activating FGF and Notch in neighboring cells. This increasingly locks the relationship between the central *atoh1a* expressing cell and surrounding, stable epithelial rosette, making it less susceptible to changes in tension along the length of the primordium. In this manner, while signaling and mechanics initiate protoneuromast formation, the self-organizing patterns they initiate are unstable and maturation of protoneuromasts initiates changes spearheaded by *atoh1a* that determine reliable, robust periodic formation of protoneuromasts and deposition of stable neuromasts by the migrating primordium.

The mechanisms that determine specification of a central sensory hair cell progenitor and reorganization of surrounding cells to form epithelial rosettes reflect the merging of two evolutionarily conserved patterning systems. The first is related to the mechanism described earlier in Drosophila development, where Notch-mediated lateral inhibition leads to the specification of a central progenitor cell within a proneural cluster (Muskavitch 1994) . Second, the combination of signaling and mechanical interactions that determine periodic clustering of cells to form epithelial rosettes has similarities to mechanisms that determine periodic clustering of cells to form feather buds and hair follicles in the skin of birds and mammals, respectively (Dalle Nogare and Chitnis 2017, Glover, Wells et al. 2017, Shyer, Rodrigues et al. 2017, Aman, Fulbright et al. 2018).

Together, the observations presented in this study illustrate how the zebrafish posterior lateral line primordium serves as an attractive model system for understanding how the patterning of cell fate specification and tissue morphogenesis can be viewed as a carefully orchestrated phenomenon resulting from a complex interplay between signaling and mechanics. It also serves as a platform for recognizing the many ways in which conserved mechanisms have evolved and have been reused in combination to coordinate reproducible and robust processes in development.

## Materials and Methods

### Zebrafish lines, embryo manipulation, and drug treatment

Zebrafish embryos were generated by natural spawning, maintained under standard conditions (28.5°C), and staged according to Kimmel et. al (Kimmel, Ballard et al. 1995). Long-term timelapse imaging was performed on *Tg(cldnb:lyn-egfp) (Haas and Gilmour 2006);TgBAC(cxcr4b:h2a-mcherry) (Wang, Yin et al. 2018);Tg(hsp:sdf1a)* (Li, Shirabe et al. 2004) triple transgenic embryos. Embryos were heat-shocked at 37.5°C for 20 minutes. The non heat-shocked triple transgenic embryos and wildtype double transgenic (*Tg(cldnb:lyn-egfp);TgBAC(cxcr4b:h2a-mcherry)*) embryos were used as controls. For PP1 drug treatment experiments, all embryos were anesthetized with MS-222 (Sigma) mixed in 1:25 proportion in E3 medium containing 20μM PP1 and mounted in 1% low-melt agarose (NuSieve GTG) containing the same concentration of the drug. Imaging was started within 10 mins. of the end of heat-shock or the addition of the drug.

### Time-lapse microscopy, image processing and quantification

Long-term timelapse confocal images of embryos were acquired using a Nikon Ti2 inverted microscope with Yokogawa CSU-W1 spinning disk confocal and the Hamamatsu Orca Flash 4 version 3 camera with a 40xW 1.1NA objective. Expression data of *Tg(cldnb:lyn-egfp)* and *TgBAC(cxcr4b:h2a-mcherry)* were acquired with 488nm and 561nm excitation lasers respectively. Images were acquired at 5-minute intervals over a period of approximately 14-16 hours. The sample stage was manually shifted to prevent the sample from going out of the imaging frame. Acquired images were then stitched using a custom in-house macro in Fiji (Schindelin, Arganda-Carreras et al. 2012).

### Particle Image Velocimetry Analysis

Particle Image Velocimetry (PIV) Analysis was done using PIVLab (Thielicke and Sonntag 2021) in MATLAB (version: 24.2 (R2024b) Update 2) (MathWorks, Natick, MA). Images were preprocessed in Fiji prior to importing into PIVLab. A Gaussian smoothing filter (σ = 2.0) was applied to the nuclei channel of acquired time-lapse movies after background subtraction to reduce noise in the images. Binary masks of nuclei were created at each timepoint using the Yen thresholding method. Smoothed nuclei images were then subtracted from the generated nuclei masks. In a densely packed tissue, such as the zebrafish PLLp, this process led to the generation of nuclei outlines, as well as graded intensities within each nucleus, with the intention that this would eventually facilitate high resolution PIV computation. These resulting images were then imported into PIVLab, where image contrast was enhanced using the CLAHE (Contrast Limited Adaptive Histogram Equalization) algorithm with a window size of 64 pixels (Pizer 1987) implemented in PIVLab. Image correlation was done in 2 passes, matching a 150x150 px^2^ window with a 75x75 px^2^ sliding window in the 1st pass, followed by matching a 75x75 px^2^ window with a 38x38 px^2^ sliding window in the 2nd pass. The last pass was repeated until the quality slope was less than 0.025. Boundary effects were neglected during computation. The magnitudes of velocities in the direction of migration were then computed in PIVLab and exported as images. These images were then superimposed on binary nuclei masks generated as detailed earlier by utilizing the “AND” image calculator function in Fiji.

### Hybridization Chain Reaction (HCR)

dkk1b and atoh1a HCR probes were purchased from Molecular Instruments, and their protocol for “whole-mount zebrafish embryos and larvae” was followed (available at: https://www.molecularinstruments.com/hcr-rnafish-protocols). *Tg(cldnb:lyn-eGFP); TgBAC(CXCR4b:h2a-mCherry); Tg(hsp:sdf1a)* triple transgenic zebrafish embryos were heat-shocked at 37°C for 20 minutes, and then fixed at various time points (0, 2, 3, and 4 hours post-heat shock) in 4% paraformaldehyde (PFA) (Electron Microscopy Sciences, 15710), overnight at 4°C in a shaker incubator. Non-heat-shocked siblings were also fixed and stored in the same conditions. The next day, embryos were washed in PBS with 0.1% Tween (Fisher Chemical, BP337-500) (PBST) and dehydrated using a methanol-PBST gradient (25%, 50%, 75%, and 100% methanol) and stored at -20°C. On the following day, embryos (8-10 per sample) were rehydrated with a methanol/PBST gradient and transferred to PBST. They were treated with proteinase K (Millipore Sigma, 3115828001), refixed in 4% PFA and prehybridized in probe hybridization buffer (HCR Buffers, Molecular Instruments) at 37°C before incubation with dkk1b and atoh1a probes diluted in hybridization buffer (4 nM each in 500 µL) overnight at 37°C. Hairpin amplifiers were prepared (30 pmol each in HCR amplification buffer) and added to the embryos after washes with probe wash buffer (HCR™ Buffers, Molecular Instruments). Amplification was conducted overnight in the dark. Embryos were washed in SSCT and PBST and then imaged using a Nikon Ti2 inverted microscope with Yokogawa CSU-W1 spinning disk confocal, Hamamatsu Orca Flash 4 version 3 camera with a 40xW 1.1NA water immersion objective.

### Computational modeling Agent Based Models (ABMs)

Agent based modeling (ABM) was done using the NetLogo programming environment (Wilensky 1999). NetLogo allows users to define rules governing the behavior of three types of agents: *turtles*, *patches* and *links*. *Turtles* are agents that can move and can be used to represent different types (breeds) of cells. *Patches* are stationary agents resembling a grid over which *turtles* move. They can represent the extracellular environment and can be used to simulate diffusion of signaling molecules produced by cells and deposited on the *patches*. *Links* are agents that connect *turtles* and can be used to represent mechanical spring-like coupling between cells resulting from the combination of inter-cellular adhesion and apical constriction. The *links* can have turtle breed-specific parameters including a *spring-constant* (stiffness or resistance to length extension of a link), *spring-length* (length all links try to achieve by pushing or pulling on connected nodes) and *repulsion-constant* (force with which connected nodes push each other to avoid overcrowding). The Fruchterman-Reingold layout algorithm, used to model the layout of nodes in a network, integrates the influence of the spring parameters (Fruchterman and Reingold 1991).

The specific parameters used in our NetLogo models, and their ranges are detailed in Supplementary Table 1. Two “breeds” of turtles were defined, **WNTers** and **FGFers,** to represent cells in a leading Wnt active and trailing FGF active domains, respectively. A third breed called **Depositers** was used in simulations described later, where progressive shrinking of the Wnt system and the periodic deposition of the cell clusters was incorporated. Various parameters could be set to determine the behavior of **WNTers** and **FGFers**. This includes their migration speed, proliferation rate and various parameters associated with *links* made with neighboring turtles. Among these parameters are the radius within which **WNTers** and **FGFers** could form *links* with neighbors, as well as the spring parameters of the *links*. In addition, the *links* were broken if they were stretched beyond a defined break-threshold and were turned over at a low but defined rate.

At each iteration of the program, **WNTers** and **FGFers** move forward a distance defined by their speed. Second, *links* are broken at a rate corresponding to their rate of turnover or if their length exceeds their break-threshold. Finally, *links* are maintained or re-made with turtles within the radius defined for that breed of *turtle.* During the simulation, the organization and movement of turtles in the primordium can be visualized in two ways; the turtles can be visualized by distinct breed color (**WNTers** - Green, **FGFers** - Yellow and **Depositers** - Red), or in shades of red reflecting the density of turtles. This “density” mode makes it easier to visualize patterns of spontaneous clustering of *turtles* under various conditions. In this mode, when turtle packing exceeds some defined density, the turtles are given a cyan color to highlight the pattern of cell clustering in the model primordium (Figure 3A).

### Cellular Potts Models (CPMs)

We implemented Cellular Potts models (CPMs) in the CompuCell3D software package (Swat, M. et al, 2012). The CPM framework allows one to simulate changes in cell morphologies based on energy minimization. Total system energy *J* is calculated as:

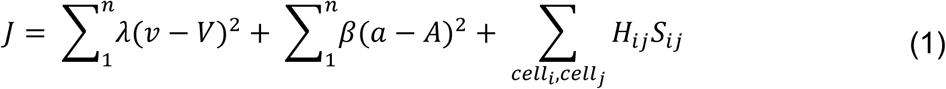

where 𝜆 is the degree of penalty imposed if the volume of the system (𝑣) varies from the target volume (𝑉). Similarly, 𝛽 is the degree of penalty imposed for variation of system area (𝑎) from target area (𝐴). 𝐻*_ij_* and 𝑆*_ij_*, where the subscripts 𝑖𝑗 refer to 𝑐𝑒𝑙𝑙*_i_* and 𝑐𝑒𝑙𝑙*_j_*, together represent cell-cell adhesion energy. Change in energy (Δ𝐽) over a Monte Carlo step can be represented as:

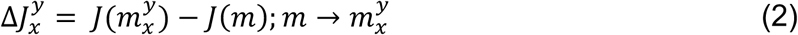

Since Δ𝐽 in the CPM framework is translated into a pixel copy, thereby effecting change in system shape, the probability of accepting such a pixel copy is determined by:

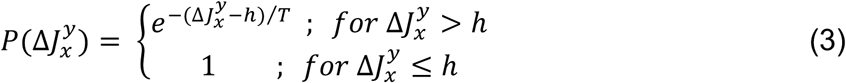

Where ℎ represents an energy threshold and *T* represents fluctuation temperature, a parameter governing stochastic dynamics of the system. Therefore, the probability of accepting a pixel copy increases with a decrease or an increase in 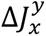 or *T* respectively.

We created a two-dimensional simulation environment (800 pixels x 300 pixels x 1 pixel) square lattice with periodic boundary conditions in the x-dimension to prevent the unintended effects of artificial rigidity and adhesivity in the system that might occur due to a fixed boundary (Figure 5B). The 2D Cellular Potts primordium model was developed representing the primordium in a side-view (Figure 5A), migrating between **skin** cells (light green) above and a layer of cells representing a thin layer of extracellular matrix (**ECM**) (brown) below, as has been recently characterized by (Yamaguchi, Zhang et al. 2022). In addition, below the **ECM** cells is a layer of cells representing a combination of the deeper **myoseptum and muscle** cells (white). Here again, the muscle layer was made thick enough (5 cells thick vertically), to minimize artificial rigidity effects of the bottom system boundary on the model primordium. The primordium itself is made of three types of cells, a) a leading domain of **Wnt** cells (blue), represents cells with a relatively non-apicobasally polarized morphology that migrate in response to a self-generated chemokine gradient, b) a trailing set of FGF cells (composite of red = apical (**fgfapi**), green = lateral (**fgflat**), and pink = basal (**fgfbas**) domains), represents polarized epithelial cells with apicobasal polarity that apically constrict and move in response to FGF signals from the leading domain, and c) a set of **sheath** cells (yellow) between the deeper **Wnt** and FGF cells and below the overlying **skin** cells. Previous studies have shown these superficial cells are essential for effective collective migration of the primordium in confinement between skin above and ECM below (Dalle Nogare, Natesh et al. 2020). An additional cell type called **Medium**, a mandatory default cell-type in CompuCell3D, signifies all the remaining space in the system, is also included in the model.

*Volume* and *Surface* plugins were used to specify 2D surface area and perimeter constraints respectively for each cell type. The *Contact* plugin helped us specify the relative adhesive energies between the various cell types. Cells that have relatively high contact energy defined between them are less likely to adhere to each other than cells that have low contact energy. Additionally, the internal adhesive energies between the three compartments of FGF cells were defined using the *ContactInternal* plugin.

Migratory forces (using the *lambdaVecX* property within the *ExternalPotential* plugin) were imparted only to **Wnt** and **sheath** cells and **fgfbas** domains of FGF cells to simulate chemokine-dependent migration of Wnt cells and the influence of FGF signaling on sheath cells and the basal feet of FGF cells. A negative *lambdaVecX* value represents the amount of force applied on the cell along the positive x-direction.

The *FocalPointPlasticity* plugin in CompuCell3D was used to create intercellular spring-like links between neighboring cells of the following types: **skin-skin**, **Wnt-Wnt**, **fgflat- fgflat**, **fgfapi-fgfapi**, and **ECM-ECM**. To maintain cohesive migration of the model primordium as a single unit, the last **Wnt** cell and the first **fgflat** cell from the leading end were connected by a spring-like link. This plugin was also used to generate intracellular links within FGF cells linking the three domains to each other, thereby simulating cytoskeletal elements within cells that impart structural rigidity and apicobasal elongation. Three parameters must be defined to simulate a spring-like link, namely: strength (equivalent to spring constant, λ), target distance (the length a link tries to attain, L), and maximum distance (the maximum length a link can stretch to before it breaks). Tuning these three parameters helped us achieve varied behaviors such as cell-scale or tissue-scale contractility, cell-cell adhesivity, cohesive collective migration, and structural rigidity. Incorporation of these links changed Eq. (1) into:

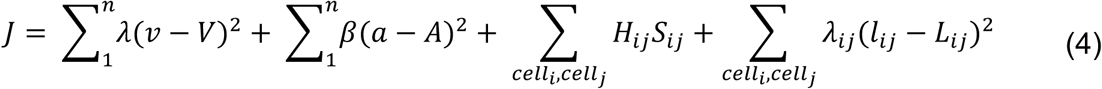

Once intercellular links between neighboring **fgfapi** compartments extended beyond their maximum length, they broke and could only be re-established if those neighboring compartments came into contact again over time. Such dynamic remodeling of the **fgfapi-fgfapi** contractile links allowed us to simulate cell-cell interactions and rosette formation in the FGF domain. The *NeighborTracker* plugin is a helper module for the *FocalPointPlasticity* plugin and was used to track all the neighbors of every cell in the system. All the parameters and variables in the model used in the current study are detailed in Supplementary Tables 2-5 (Appendix). Additionally, the fluctuation temperature *T* was set to 20 and neighbor order, governing the region of influence of a cell on its neighbors, was set to 3.

## Acknowledgements

The authors would like to thank Julia Rashid and Seth Entriken for help with editing the manuscript.

## Author Contributions

Conceptualization: AB Chitnis, A Mukherjee; Methodology: AB Chitnis, A Mukherjee, M Hilzendeger, S Fatma, D Dalle Nogare; Software: AB Chitnis, A Mukherjee; Validation: AB Chitnis, A Mukherjee, A Rinvelt, M Schupp; Formal analysis: A Mukherjee, M Hilzendeger; Investigation: A Mukherjee, M Hilzendeger, S Fatma; Resources: AB Chitnis; Data curation: A Mukherjee, A Rinvelt, AB Chitnis; Writing - original draft: AB Chitnis, A Mukherjee; Writing - review & editing: AB Chitnis, A Mukherjee; Visualization: AB Chitnis, A Mukherjee, M Hilzendeger, A Rinvelt; Supervision: AB Chitnis ; Project administration: AB Chitnis; Funding acquisition: AB Chitnis

## Code and Data Availability

All the data generated for this study are available upon request. All requests should be addressed to Ajay Chitnis (email). Codes are available at <u>GitHub -</u> <u>abhimukherjee1/models_for_mechanics_paper</u>

## Supplementary Figure Legends

**Figure S1:**
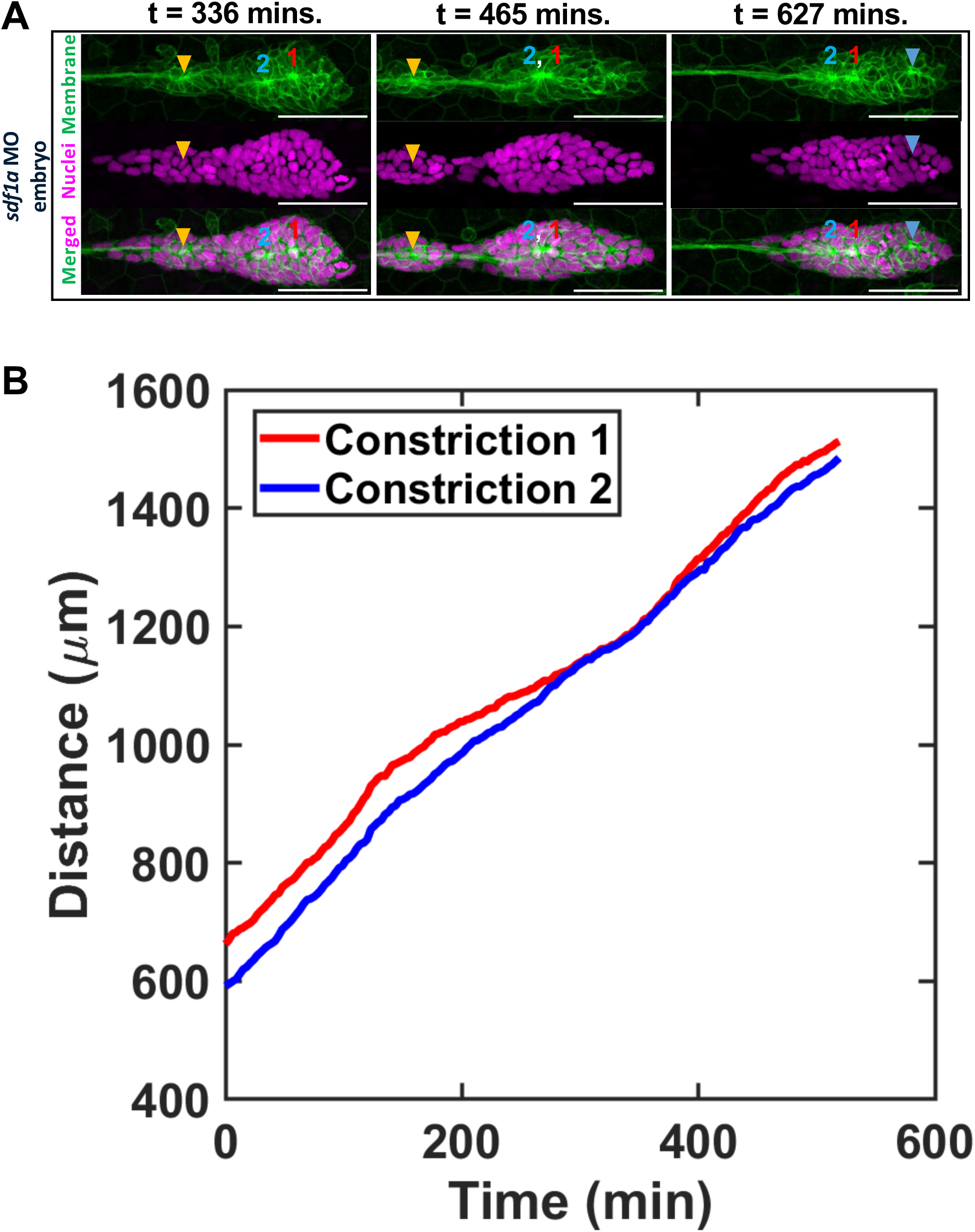
**(A)** Panels showing rosettes fusing and splitting in an *sdf1a* morpholino-injected embryo. Membrane and nuclei are shown in green and magenta respectively. The constrictions of interest are marked by 1 (red) and 2 (blue). A stable depositing rosette is marked by an orange arrowhead, and a nascent protoneuromast that forms after constrictions 1 and 2 fuse is marked by a light blue arrowhead. **(B)** Tracking constrictions 1 and 2 over time shows them fusing and splitting up. Tracking starts when constriction 2 forms. Note that this leads to a slight shift in time in the panels **(A)** and **(B)** (Time = 0 in panel **(B)** corresponds to 126 mins. in panel **(A)**).

**Figure S2:**
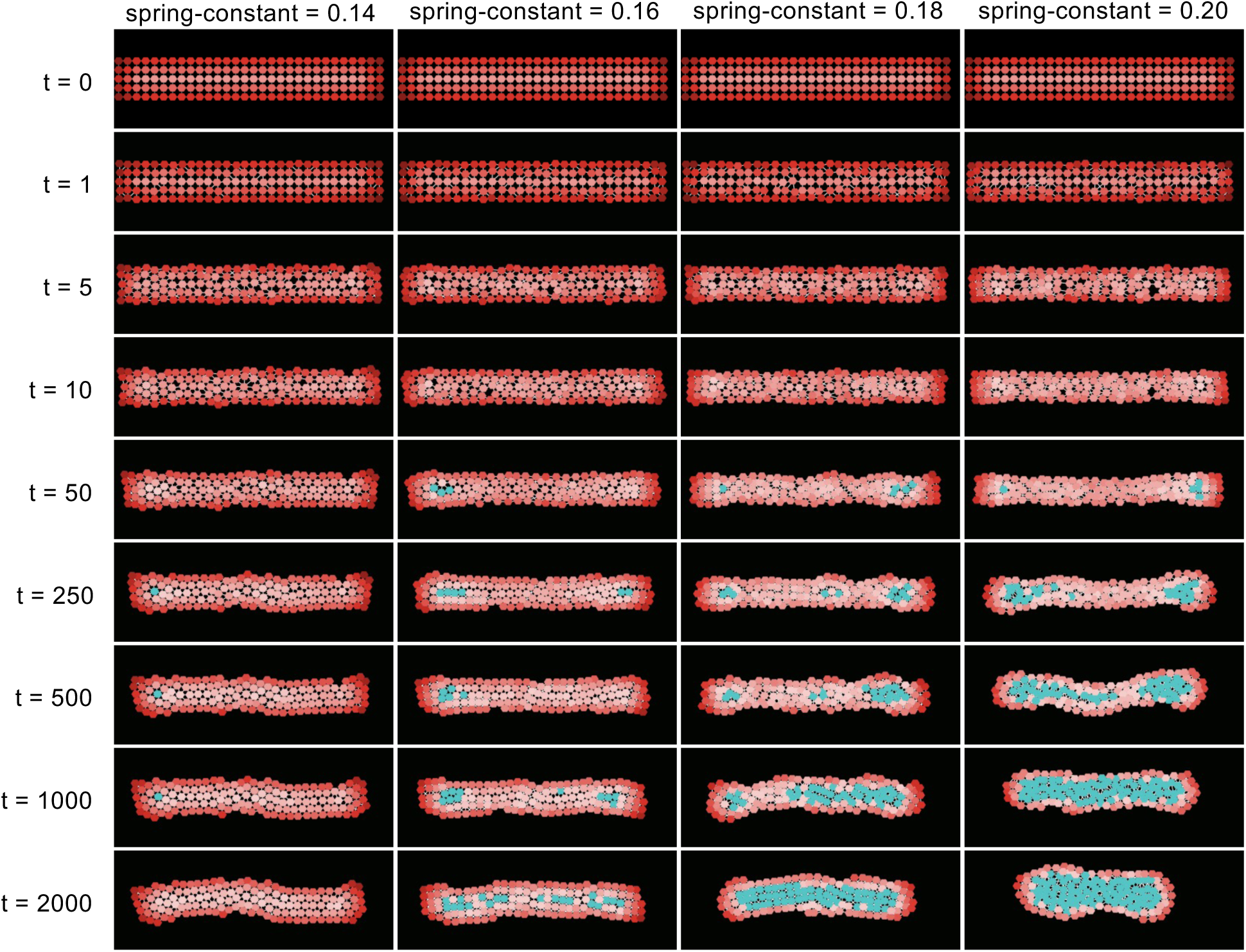
Evolution of turtle aggregation dynamics over time as a function of spring-constant of links.

**Figure S3:**
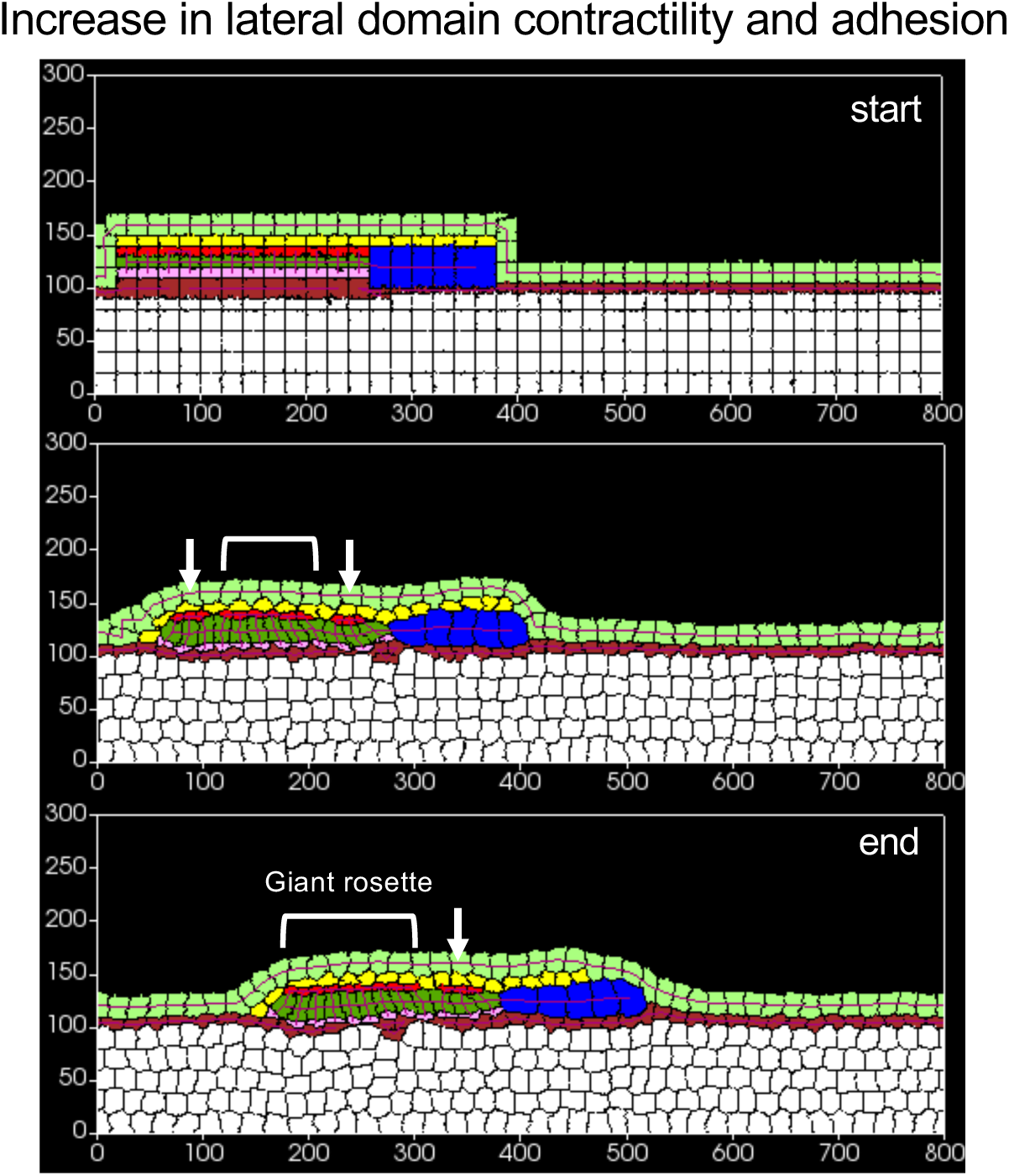
Example simulation where lateral domain contractility (𝜆*_fgflat−fgflat_* = 40) and cell-cell adhesion (𝐸*_fgflat−fgflat_* = 1) have been increased. Panels depict start (top), an interim timestep (middle), and end (bottom) of simulation. Relatively small rosettes are marked with arrows, and relatively larger ones are marked with brackets.

## Appendix

### Supplementary Tables

**Table 1:** Details of the agent-based model created in NetLogo.

| Variable Name | Physiological representation | Value |
| --- | --- | --- |
| Wnter speed | Speed of leading cells | 0, 0.015 – 0.018 |
| FGFer speed | Speed of trailing cells | 0, 0.015 – 0.018 |
| PLLp width | PLLp width | 5 |
| PLLp length | PLLp length | 30 |
| Proliferation rate | Rate of cell divisions | $5 \times 10^{-4}$ |
| Wnt fraction | Fraction of PLLp with high Wnt activity | 0.6 or 1 |
| Wnt size |  | 30 |
| FGF size |  | 30 |
| Wnt spring constant | Adhesivity + Contractility of leading cells | 0.14 – 0.2 |
| FGF spring constant | Adhesivity + Contractility of trailing cells | 0.14 – 0.2 |
| Wnt spring length | Refers to the region of influence of intercellular adhesion in the leading and trailing domains | 0.1 |
| FGF spring length |  | 0.1 |
| Wnt repulsion constant | Refers to how close cells are allowed to come before being repelled | 0.03 |
| FGF repulsion constant |  | 0.03 |
| Wnt radius |  | 1.25 |
| FGF radius |  | 1.25 |
| Cluster threshold | For visualization purposes only | 32 |
| Density scale | For visualization purposes only | 36 |

**Table 2:**
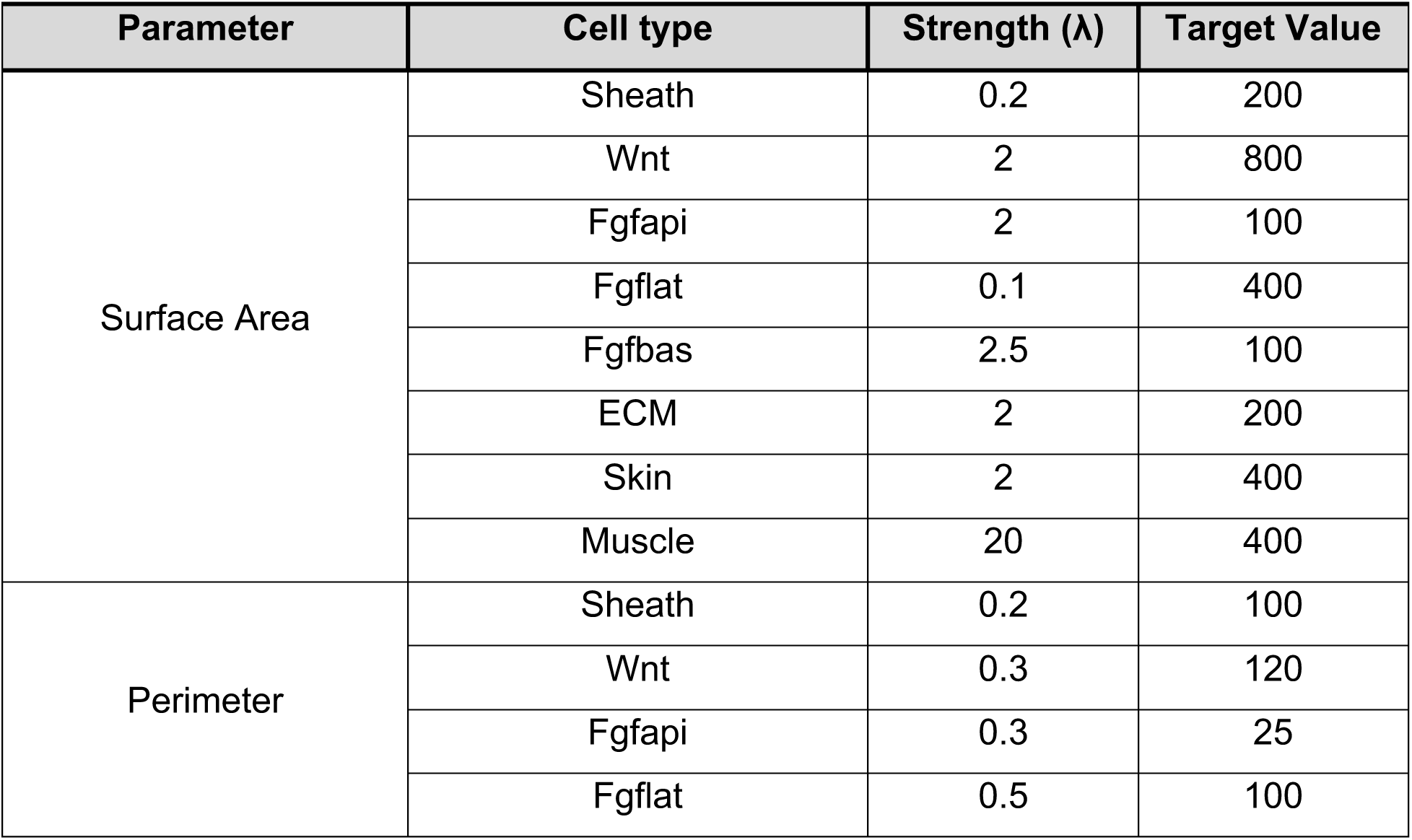
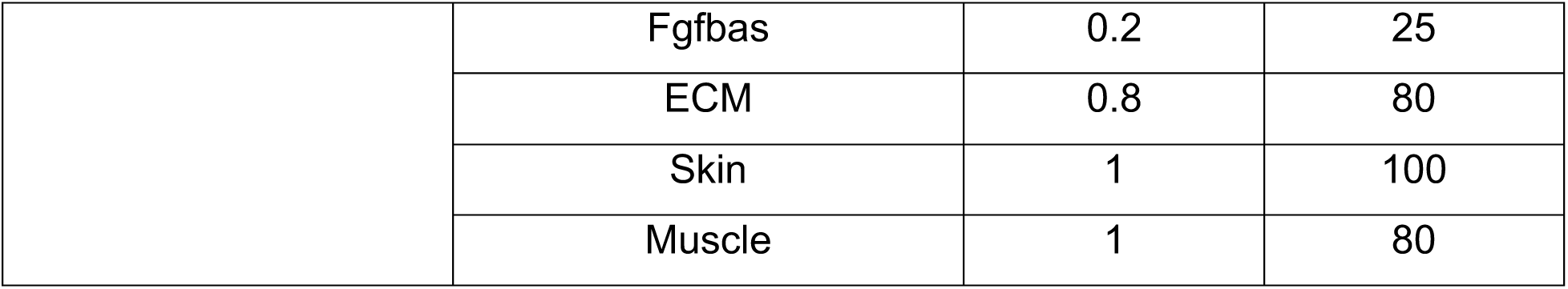
Cell shape constraints of the CPM created in CompuCell3D.

| Parameter | Cell type | Strength ($\lambda$) | Target Value |
| --- | --- | --- | --- |
| Surface Area | Sheath | 0.2 | 200 |
|  | Wnt | 2 | 800 |
|  | Fgfapi | 2 | 100 |
|  | Fgflat | 0.1 | 400 |
|  | Fgfbas | 2.5 | 100 |
|  | ECM | 2 | 200 |
|  | Skin | 2 | 400 |
|  | Muscle | 20 | 400 |
| Perimeter | Sheath | 0.2 | 100 |
|  | Wnt | 0.3 | 120 |
|  | Fgfapi | 0.3 | 25 |
|  | Fgflat | 0.5 | 100 |
|  | Fgfbas | 0.2 | 25 |
|  | ECM | 0.8 | 80 |
|  | Skin | 1 | 100 |
|  | Muscle | 1 | 80 |

**Table 3:** Cell-cell adhesion energies within the CPM.

| Parameter | Cell-Cell interaction | Contact Energy |
| --- | --- | --- |
| Contact Energies | Wnt-Sheath | 6 |
|  | Wnt-Wnt | 1 |
|  | Wnt-Fgfapi | 3 |
|  | Wnt-Fgflat | 0.5 |
|  | Wnt-Fgfbas | 3 |
|  | Wnt-ECM | 5 |
|  | Wnt-Skin | 5 |
|  | Wnt-Muscle | 10 |
|  | Fgfapi-Sheath | 6 |
|  | Fgfapi-Fgfapi | 1 |
|  | Fgfapi-Fgflat | 10 |
|  | Fgfapi-Fgfbas | 20 |
|  | Fgfapi-ECM | 20 |
|  | Fgfapi-Skin | 20 |
|  | Fgfapi-Muscle | 20 |
|  | Fgflat-Sheath | 10 |
|  | Fgflat-Fgflat | [1, 5] |
|  | Fgflat-Fgfbas | 10 |
|  | Fgflat-ECM | 20 |
|  | Fgflat-Skin | 20 |
|  | Fgflat-Muscle | 20 |
|  | Fgfbas-Sheath | 5 |
|  | Fgfbas-Fgfbas | 8 |
|  | Fgfbas-ECM | 2 |
|  | Fgfbas-Skin | 20 |
|  | Fgfbas-Muscle | 10 |
|  | ECM-Sheath | 8 |
|  | ECM-ECM | 0.5 |
|  | ECM-Skin | 6 |
|  | ECM-Muscle | 1 |
|  | Sheath-Sheath | 6 |
|  | Sheath-Skin | 4 |
|  | Sheath-Muscle | 20 |
|  | Skin-Skin | 0 |
|  | Skin-Muscle | 10 |
|  | Muscle-Muscle | 1 |
| Internal Contact Energies<br>(only for compartmental<br>FGF cells) | Fgflat-Fgfbas | 1 |
|  | Fgfapi-Fgflat | 1 |
|  | Fgfapi-Fgfbas | 10 |

**Table 4:** Cell Migration details in CPM.

| Cell type | Migratory Force ($\lambda_x$) (+x-direction) | Cell fluctuation amplitude |
| --- | --- | --- |
| Wnt | [-0.1, -2] | [10, 50] |
| Fgfbas | -3 | 30 |
| Sheath | -1 | 50 |

**Table 5:** *FocalPointPlasticity* links details in CPM.

| Cell-cell interaction | Strength ($\lambda$) | Target distance | Maximum distance |
| --- | --- | --- | --- |
| Wnt-Wnt | 5 | 10 | 200 |
| Wnt-Fgflat | 10 | 30 | 40 |
| Fgfapi-Fgfapi | 5 | 8 | 20 |
| Fgflat-Fgflat | [10, 40] | 8 | 100 |
| Skin-Skin | 10 | 15 | 100 |
| ECM-ECM | 4 | 10 | 40 |
| Fgfapi-Fgflat | 8 | 15 | 30 |
| Fgflat-Fgfbas | 8 | 15 | 30 |

## Supplementary Movies

**Movie S1:**

Movie of a migrating wildtype control PLLp showing cell membranes (top), nuclei (middle), and merged (bottom) images.

**Movie S2:**

Movie of a migrating PLLp in a heat-shocked *Tg(hsp:sdf1a)* embryo showing cell membranes (top), nuclei (middle), and merged (bottom) images.

**Movie S3:**

Movie of the migrating wildtype control PLLp from Movie S1, showing cell membranes (top) and the corresponding PIV colormap overlaid on cell nuclei (bottom).

**Movie S4:**

Movie of a migrating PLLp in the heat-shocked *Tg(hsp:sdf1a)* embryo from Movie S2, showing cell membranes (top) and the corresponding PIV colormap overlaid on cell nuclei (bottom).

**Movie S5:**

Movie of a migrating PLLp in a standard control morpholino-injected embryo showing cell membranes (top), nuclei (middle), and merged (bottom) images.

**Movie S6:**

Movie of a migrating PLLp in a 2ng sdf1a morpholino-injected embryo showing cell membranes (top), nuclei (middle), and merged (bottom) images.

**Movie S7:**

Movie of a migrating PLLp treated with 20μM PP1 showing cell membranes (top) and the corresponding PIV colormap overlaid on cell nuclei (bottom).

**Movie S8:**

Movie of a representative agent-based model simulation. Simulation showing aggregation of turtles without migration (Top), with **WNTers** moving faster than **FGFers** (middle), and with **WNTers** moving slower than **FGFers** (bottom).

**Movie S9:**

Movie of a representative agent-based model simulation showing the shrinkage of **WNTers** and deposition of **Depositers**

**Movie S10:**

Movie of a representative Cellular Potts model PLLp simulating WT control conditions.

**Movie S11:**

Movie of a representative Cellular Potts model PLLp where leading cells slow down before resuming migration, thereby simulating the heat-shocked *Tg(hsp:sdf1a)* phenotype.

**Movie S12:**

Movie of a representative Cellular Potts model PLLp where an increase in lateral domain contractility and adhesion leads to formation of a large rosette.

## References

Aman, A. and T. Piotrowski (2008). "Wnt/beta-catenin and Fgf signaling control collective cell migration by restricting chemokine receptor expression." Dev Cell 15(5): 749–761.

Aman, A. J., A. N. Fulbright and D. M. Parichy (2018). "Wnt/beta-catenin regulates an ancient signaling network during zebrafish scale development." Elife 7.

Bussmann, J. and E. Raz (2015). "Chemokine-guided cell migration and motility in zebrafish development." EMBO J 34(10): 1309–1318.

Dalle Nogare, D. and A. B. Chitnis (2017). "A framework for understanding morphogenesis and migration of the zebrafish posterior Lateral Line primordium." Mech Dev 148: 69–78.

Dalle Nogare, D. and A. B. Chitnis (2017). "Self-organizing spots get under your skin." PLoS Biol 15(12): e2004412.

Dalle Nogare, D. and A. B. Chitnis (2020). "NetLogo agent-based models as tools for understanding the self-organization of cell fate, morphogenesis and collective migration of the zebrafish posterior Lateral Line primordium." Semin Cell Dev Biol 100: 186–198.

Dalle Nogare, D., K. Somers, S. Rao, M. Matsuda, M. Reichman-Fried, E. Raz and A. B. Chitnis (2014). "Leading and trailing cells cooperate in collective migration of the zebrafish posterior lateral line primordium." Development 141(16): 3188–3196.

Dalle Nogare, D. E., N. Natesh, H. D. Vishwasrao, H. Shroff and A. B. Chitnis (2020). "Zebrafish Posterior Lateral Line primordium migration requires interactions between a superficial sheath of motile cells and the skin." Elife 9.

Dambly-Chaudiere, C., N. Cubedo and A. Ghysen (2007). "Control of cell migration in the development of the posterior lateral line: antagonistic interactions between the chemokine receptors CXCR4 and CXCR7/RDC1." BMC Dev Biol 7: 23.

Dona, E., J. D. Barry, G. Valentin, C. Quirin, A. Khmelinskii, A. Kunze, S. Durdu, L. R. Newton, A. Fernandez-Minan, W. Huber, M. Knop and D. Gilmour (2013). "Directional tissue migration through a self-generated chemokine gradient." Nature 503(7475): 285–289.

Ernst, S., K. Liu, S. Agarwala, N. Moratscheck, M. E. Avci, D. Dalle Nogare, A. B. Chitnis, O. Ronneberger and V. Lecaudey (2012). "Shroom3 is required downstream of FGF signalling to mediate proneuromast assembly in zebrafish." Development 139(24): 4571–4581.

Fruchterman, T. M. J. and E. M. Reingold (1991). "Graph Drawing by Force-Directed Placement." Software-Practice & Experience 21(11): 1129–1164.

Ghysen, A. and C. Dambly-Chaudiere (2007). "The lateral line microcosmos." Genes Dev 21(17): 2118–2130.

Gierer, A. and H. Meinhardt (1972). "A theory of biological pattern formation." Kybernetik 12(1): 30–39.

Glover, J. D., K. L. Wells, F. Matthaus, K. J. Painter, W. Ho, J. Riddell, J. A. Johansson, M. J. Ford, C. A. B. Jahoda, V. Klika, R. L. Mort and D. J. Headon (2017). "Hierarchical patterning modes orchestrate hair follicle morphogenesis." PLoS Biol 15(7): e2002117.

Haas, P. and D. Gilmour (2006). "Chemokine signaling mediates self-organizing tissue migration in the zebrafish lateral line." Dev Cell 10(5): 673–680.

Harris, A. K., D. Stopak and P. Warner (1984). "Generation of spatially periodic patterns by a mechanical instability: a mechanical alternative to the Turing model." J Embryol Exp Morphol 80: 1–20.

Kimmel, C. B., W. W. Ballard, S. R. Kimmel, B. Ullmann and T. F. Schilling (1995). "Stages of embryonic development of the zebrafish." Dev Dyn 203(3): 253–310.

Kozlovskaja-Gumbriene, A., R. Yi, R. Alexander, A. Aman, R. Jiskra, D. Nagelberg, H. Knaut, M. McClain and T. Piotrowski (2017). "Proliferation-independent regulation of organ size by Fgf/Notch signaling." Elife 6.

Lau, S., A. Feitzinger, G. Venkiteswaran, J. Wang, S. W. Lewellis, C. A. Koplinski, F. C. Peterson, B. F. Volkman, M. Meier-Schellersheim and H. Knaut (2020). "A negative-feedback loop maintains optimal chemokine concentrations for directional cell migration." Nat Cell Biol 22(3): 266–273.

Lecaudey, V., G. Cakan-Akdogan, W. H. Norton and D. Gilmour (2008). "Dynamic Fgf signaling couples morphogenesis and migration in the zebrafish lateral line primordium." Development 135(16): 2695–2705.

Li, Q., K. Shirabe and J. Y. Kuwada (2004). "Chemokine signaling regulates sensory cell migration in zebrafish." Dev Biol 269(1): 123–136.

Matsuda, M. and A. B. Chitnis (2010). "Atoh1a expression must be restricted by Notch signaling for effective morphogenesis of the posterior lateral line primordium in zebrafish." Development 137(20): 3477–3487.

Matsuda, M., D. D. Nogare, K. Somers, K. Martin, C. Wang and A. B. Chitnis (2013). "Lef1 regulates Dusp6 to influence neuromast formation and spacing in the zebrafish posterior lateral line primordium." Development 140(11): 2387–2397.

Meinhardt, H. (2012). "Turing’s theory of morphogenesis of 1952 and the subsequent discovery of the crucial role of local self-enhancement and long-range inhibition." Interface Focus 2(4): 407–416.

Muskavitch, M. A. (1994). "Delta-notch signaling and Drosophila cell fate choice." Dev Biol 166(2): 415–430.

Neelathi, U. M., D. Dalle Nogare and A. B. Chitnis (2018). "Cxcl12a induces snail1b expression to initiate collective migration and sequential Fgf-dependent neuromast formation in the zebrafish posterior lateral line primordium." Development 145(14).

Nogare, D. D., M. Nikaido, K. Somers, J. Head, T. Piotrowski and A. B. Chitnis (2017). "In toto imaging of the migrating Zebrafish lateral line primordium at single cell resolution." Dev Biol 422(1): 14–23.

Oster, G. F., J. D. Murray and A. K. Harris (1983). "Mechanical aspects of mesenchymal morphogenesis." J Embryol Exp Morphol 78: 83–125.

Pizer, S. M., Amburn, E. P., Austin, J. D., Cromartie, R., Geselowitz, A., Greer, T. (1987). "Adaptive histogram equalization and its variations." Computer Vision Graphics and Image Processing 39(3): 13.

Sapede, D., M. Rossel, C. Dambly-Chaudiere and A. Ghysen (2005). "Role of SDF1 chemokine in the development of lateral line efferent and facial motor neurons." Proc Natl Acad Sci U S A 102(5): 1714–1718.

Schindelin, J., I. Arganda-Carreras, E. Frise, V. Kaynig, M. Longair, T. Pietzsch, S. Preibisch, C. Rueden, S. Saalfeld, B. Schmid, J. Y. Tinevez, D. J. White, V. Hartenstein, K. Eliceiri, P. Tomancak and A. Cardona (2012). "Fiji: an open-source platform for biological-image analysis." Nat Methods 9(7): 676–682.

Shyer, A. E., A. R. Rodrigues, G. G. Schroeder, E. Kassianidou, S. Kumar and R. M. Harland (2017). "Emergent cellular self-organization and mechanosensation initiate follicle pattern in the avian skin." Science 357(6353): 811–815.

Thielicke, W. and R. Sonntag (2021). "Particle Image Velocimetry for MATLAB: Accuracy and enhanced algorithms in PIVlab." Journal of Open Research Software.

Turing, A. M. (1990). "The chemical basis of morphogenesis. 1953." Bull Math Biol 52(1-2): 153–197; discussion 119-152.

Valentin, G., P. Haas and D. Gilmour (2007). "The chemokine SDF1a coordinates tissue migration through the spatially restricted activation of Cxcr7 and Cxcr4b." Curr Biol 17(12): 1026–1031.

Venkiteswaran, G., S. W. Lewellis, J. Wang, E. Reynolds, C. Nicholson and H. Knaut (2013). "Generation and dynamics of an endogenous, self-generated signaling gradient across a migrating tissue." Cell 155(3): 674–687.

Wang, J., Y. Yin, S. Lau, J. Sankaran, E. Rothenberg, T. Wohland, M. Meier-Schellersheim and H. Knaut (2018). "Anosmin1 Shuttles Fgf to Facilitate Its Diffusion, Increase Its Local Concentration, and Induce Sensory Organs." Dev Cell 46(6): 751–766 e712.

Wilensky, U. (1999). "Netlogo." from http://ccl.northwestern.edu/netlogo/.

Wong, M., L. R. Newton, J. Hartmann, M. L. Hennrich, M. Wachsmuth, P. Ronchi, A. Guzman-Herrera, Y. Schwab, A. C. Gavin and D. Gilmour (2020). "Dynamic Buffering of Extracellular Chemokine by a Dedicated Scavenger Pathway Enables Robust Adaptation during Directed Tissue Migration." Dev Cell 52(4): 492–508 e410.

Yamaguchi, N., Z. Zhang, T. Schneider, B. Wang, D. Panozzo and H. Knaut (2022). "Rear traction forces drive adherent tissue migration in vivo." Nat Cell Biol 24(2): 194–204.

